# Repurposing tofacitinib as an anti-myeloma therapeutic to reverse growth-promoting effects of the bone marrow microenvironment

**DOI:** 10.1101/143206

**Authors:** Christine Lam, Megan Murnane, Hui Liu, Geoffrey A. Smith, Sandy Wong, Jack Taunton, Jun O. Liu, Constantine S. Mitsiades, Byron C. Hann, Blake T. Aftab, Arun P. Wiita

## Abstract

The myeloma bone marrow microenvironment promotes proliferation of malignant plasma cells and resistance to therapy. Interleukin-6 (IL-6) and downstream JAK/STAT signaling are thought to be central components of these microenvironment-induced phenotypes. In a prior drug repurposing screen, we identified tofacitinib, a pan-JAK inhibitor FDA-approved for rheumatoid arthritis, as an agent that may reverse the tumor-stimulating effects of bone marrow mesenchymal stromal cells. Here, we validated both *in vitro*, in stromal-responsive human myeloma cell lines, and *in vivo*, in orthotopic disseminated murine xenograft models of myeloma, that tofacitinib showed both single-agent and combination therapeutic efficacy in myeloma models. Surprisingly, we found that ruxolitinib, an FDA-approved agent targeting JAK1 and JAK2, did not lead to the same anti-myeloma effects. Combination with a novel irreversible JAK3-selective inhibitor also did not enhance ruxolitinib effects. RNA-seq and unbiased phosphoproteomics revealed that marrow stromal cells stimulate a JAK/STAT-mediated proliferative program in myeloma plasma cells, and tofacitinib reversed the large majority of these pro-growth signals. Taken together, our results suggest that tofacitinib specifically reverses the growth-promoting effects of the tumor microenvironment through blocking an IL-6-mediated signaling axis. As tofacitinib is already FDA-approved, these results can be rapidly translated into potential clinical benefits for myeloma patients.

## Introduction

Multiple myeloma (MM) is the second-most common hematologic malignancy in the United States and still has no known cure. A major therapeutic challenge in MM is that patients undergo numerous cycles of response to therapy followed by disease relapse after the development of resistance. Years of research have revealed that a major driver of malignant plasma cell proliferation, as well as therapeutic resistance, is signaling to the tumor cells from the bone marrow microenvironment (reviewed in refs. 1-3). Cell types within the bone marrow that influence myeloma plasma cells include mesenchymal stromal cells, osteoblasts, osteoclasts, and multiple classes of immune cells^1–3^. Overcoming the growth-promoting phenotype of the bone marrow microenvironment is thought to be a promising therapeutic strategy in MM. One approach to identifying new therapeutic agents for many diseases is drug repurposing.

In this context, a large library of drugs, all of which are either FDA-approved or at the minimum shown to be safe in humans, is screened against the biological system of interest^4,5^. The premise behind these screens is that small molecules initially designed for one indication may actually have beneficial effects across other diseases. In fact the use of thalidomide in multiple myeloma is one of the most impactful examples of successful drug repurposing. If new indications are found for already existing drugs, clinical development times and associated costs can be drastically reduced, accelerating potential benefits to patients^6,7^.

To identify agents which may reverse the tumor-promoting effects of the MM bone marrow microenvironment, we recently reported a repurposing screen of 2,684 compounds, against three MM cell lines, either grown alone (monoculture) or in co-culture with MM patient-derived bone marrow mesenchymal stromal cells^8^. From that screen, we identified tofacitinib citrate, an FDA-approved small molecule for the treatment of rheumatoid arthritis (RA), sold under the trade name Xeljanz, as an agent which may reverse stromal-induced growth proliferation of malignant plasma cells.

Tofacitinib citrate is a potent inhibitor of all four members of the Janus kinase (JAK) family, with preferential inhibition of JAK1 and JAK3 over JAK2 and TYK2 in cellular assays^9^. JAK signaling, mediated by the downstream STAT transcription factors, is necessary for lymphocyte stimulation in response to encountered antigens^10^. Therefore, JAK inhibition holds promise for the treatment of autoimmune diseases like RA^11^. In parallel, the JAKs have gained interest as therapeutic targets in MM as they mediate signaling via interleukin-6 (IL-6). IL-6 is secreted by many cell types within the bone marrow microenvironment, as well as by malignant plasma cells themselves, and it is thought that proliferation of malignant plasma cells within the human bone marrow is dependent on this cytokine^12^. This dependence on IL-6 was underscored by the recent development of a patient-derived xenograft model of MM, where primary plasma cell growth only occurred in immunocompromised mouse bone marrow after knock-in of human IL-6^13^.

Therefore, abrogating IL-6 signaling via JAK inhibition is a promising therapeutic strategy in MM^14^. In fact, a number of groups have aimed to target IL-6 stimulation via novel small molecule inhibitors that variously target JAK1/JAK2 (ref. 15,16), JAK2 (ref. 17–19), or all four JAKs (ref. 20), with reported preclinical therapeutic efficacy in MM. However, per the registry at clinicaltrials.gov, none of these experimental JAK inhibitors have ever entered into clinical trials with MM as an indication. Therefore, all of these agents are very far from use in MM patients, if they ever become available at all. Here, we demonstrate that the already FDA-approved agent tofacitinib has robust preclinical activity in MM models. We further use RNA-seq and unbiased mass spectrometry-based phosphoproteomics to delineate pro-proliferative signals from the bone marrow stroma and show that they are largely reversed by tofacitinib treatment. Furthermore, we find that an alternate repurposing candidate, the FDA-approved JAK1/2 inhibitor ruxolitinib, surprisingly does not show the same anti-myeloma properties. Therefore, our results support the rapid repurposing of tofacitinib as an anti-myeloma therapeutic, to reverse the pro-growth effects of the bone marrow microenvironment and potentiate the effects of existing myeloma therapies.

## Materials and Methods

### Cell culture conditions

HS5 and HS27A stromal cell lines were obtained from American Type Culture Collection (ATCC). MM.1S mC/Luc, stably expressing mCherry and luciferase, were generated from parental MM.1S line obtained from ATCC, as previously described^21^. The RPMI8266 mC/Luc, U266 mC/Luc, and JJN3 mC/Luc lines were generated from parental cell lines obtained from the Deutsche Sammlung von Mikroorganimen und Zellkuturen (DSMZ) repository. These cell lines were stably transduced with a lentiviral expression plasmid constitutively expressing luciferase and mCherry, generated and kindly provided by Dr. Diego Acosta-Alvear at UCSF. L363 cell line was also obtained from DSMZ. AMO-1 cells were kindly provided by Dr. Cristoph Driessen at Kantonsspital St. Gallien, Switzerland. KMS11 cells were obtained from JCRB Cell Bank. INA-6 cells were obtained from Dr. Renate Burger at University Hospital Schleswig-Holstein, Germany. All cells, including patient bone marrow mononuclear cells, were maintained in complete media with RPMI-1640 supplemented with 10% FBS (Gemini), 1% penicillin-streptomycin (UCSF), and 2 mM L-Glutamine (UCSF) with 5% CO_2_. INA-6 media was supplemented with 50 ng/mL recombinant human IL-6 (ProSpec).

### MM and BMSC coculture and viability testing

#### Dose response

Cocultures were seeded into 384 well plates (Corning) with the Multidrop Combi (Thermo Scientific). 800 stromal cells were seeded and incubated overnight. 17 hours later, 700 myeloma cells were added on top of stromal cells. On the third day, 24 hours after addition of myeloma cells, cocultures were treated with tofacitinib (LC Laboratories), ruxolitinib (Selleck Chemicals), or JAK3i (ref. 22). For drug combination studies, on the fourth day, melphalan (Sigma Aldrich) or carfilzomib (Selleck Chemicals) were additionally added to cocultures. On the fifth day, myeloma cell viability was detected with addition of luciferin (Gold Biotechnology) and read for luminescence on Glomax Explorer plate reader (Promega) as previously described^21^. For monoculture studies cell viability was measured using Cell-Titer Glo reagent (Promega). All measurements were performed in quadruplicate. All viability data are reported as normalized to DMSO-treated cell line in monoculture.

### RNA-seq

For co-culture RNA-seq, 5x10^6^ MM.1S cells were grown in co-culture with 3x10^6^ HS5 cells for 24 hr. All cells were harvested by Accutase incubation for 10 min at 37 C to separate MM.1S from HS5 into a single-cell suspension. Cells were then separated using CD138+ MicroBeads (Miltenyi) on a Miltenyi MidiMACS system. CD138+ enrichment to >95% was verified by flow cytometry for mCherry expression (Supplementary Fig. 1). MM.1S harvested from co-culture, as well as MM.1S and HS5 grown in monoculture, were then processed for mRNA-seq as previously described^23^. RNA-seq performed on HS5 stromal cells alone further allowed for filtering of highly expressed stromal cell genes that could interfere with analysis of MM.1S cells in co-culture even at <5% contamination. Significantly upregulated transcripts were identified by DESeq^24^ and bioinformatic analysis was performed using Enrichr^25^. Raw sequencing data are available at the GEO repository (Accession number GSE99293).

### Western Blot Analysis

For co-culture studies, 3x10^5^ HS5 were seeded into 6 well plate. 17 hours later, 5x10^6^ MM.1S mC/Luc cells were added on top of stromal cell layer. For monoculture studies 5x10^6^ MM1.S cells alone were used. Cells were treated with tofacitinib or ruxolitinib from 0-24 hours and the supernatant, containing all MM1.S cells in suspension, harvested for processing by centrifugation. All Western blots were performed at in biological duplicate with representative blots displayed. pSTAT3 (Tyr705), STAT3, pSTAT1 (Tyr701)(58D6), STAT1, pJAK1(Tyr1022/1023), JAK1, pJAK2 (Tyr1008), JAK2, pJAK3 (Tyr980/981), JAK3, pTYK2 (Tyr1054/1055), pSTAT5 (Tyr694), and STAT5 antibodies were purchased from Cell Signaling Technology. Tyk2 antibody was purchased from Santa Cruz Biotechnology. Western blot was carried out as previously described^23^.

### Liquid chromatography–tandem mass spectrometry phosphoproteomics

For co-culture experiments, 5x10^6^ HS5 were seeded into a T75 flask. 17 hours later, cultures were washed with PBS, before addition of 10^7^ MM.1S mC/Luc cells. 24 hours later, cocultures were treated with 1 μM tofacitinib for 1.5 hours and 24 hours. On the third day, MM1.S cells in suspension were harvested by aspiration, centrifuged, washed with PBS, and flash-frozen prior to analysis. For untreated MM1.S monoculture experiments 10^7^ cells were used. For sample preparation frozen cell pellets were lysed in 8M urea. 1 mg of total protein was then reduced in TCEP and free cysteines alkylated with iodoacetamide. Proteins were then digested at room temperature for 18 hours with trypsin. Peptides were desalted, lyophilized, and enriched for phosphopeptides using immobilized-metal affinity column (IMAC) with Fe-NTA loaded beads^26^. Phosphopeptides were analyzed on a Thermo Q-Exactive Plus mass spectrometer coupled to a Dionex Ultimate 3000 NanoRSLC liquid chromatography instrument with 3.5 hr linear gradient. Spectra were acquired in data-dependent acquisition mode at 70,000 resolution. Raw mass spectrometry data for two biological replicates were processed using Maxquant v1.5 (ref. 27) versus the Uniprot human proteome (downloaded Feb. 20, 2017; 157,537 entries), with phospho (STY) selected as a fixed modification and match between runs enabled. Only phosphopeptides with measured MaxQuant MS1 intensity in all samples were used for analysis. Median intensity normalization of all phosphopeptides in the sample was performed prior to analysis. Raw proteomic data files are available at the ProteomXchange PRIDE repository (Accession number PXD006581).

### Xenograft mouse model

NOD.*Cg-Prkdc^scid^Il2rg^tm1Wjl^*/SzJ (NSG) mice were obtained from Jackson laboratory. 10^6^ MM.1S mC/Luc or U266 mC/Luc, stably expressing luciferase, were transplanted via tail vein injection into each mouse. Tumor burden was assessed through weekly bioluminescent imaging, beginning 13 days after implantation and same day as treatment initiation. Mice were treated for 4 weeks with vehicle, tofacitinib, carfilzomib, and combination of tofacitinib and carfilzomib as indicated (5 mice/arm.) Tofacitinib was formulated in 50% DMSO, 10% PEG 400, and 40% water and administered at 21.5 mg/kg daily by continuous subcutaneous infusion. Carfilzomib was formulated in 10% cyclodextrin (Captisol) and 10 mM sodium citrate, pH 3.5, administered at 2 mg/kg 2x/week IV. Vehicle was carfilzomib formulation administered 2x/week IV. All mouse studies were performed according to UCSF Institutional Animal Care and Use Committee-approved protocols.

### Patient samples

De-identified primary MM bone marrow samples were obtained from the UCSF Hematologic Malignancy Tissue Bank in accordance with UCSF Committee on Human Research-approved protocols. Bone marrow mononuclear cells were isolated by density gradient centrifugation Histopaque-1077 (Sigma Aldrich), then adjusted to 2 x 10^5^/well in a 96 well plate. Primary cells were stimulated with 50 ng/ml recombinant human IL-6 (ProsPec) for 17 hours before treatment with tofacitinib for 24 hours. Cells were then stained with Alexa-Fluor 647 mouse anti-human CD138 antibody (BD Pharmingen), Alexa-Fluor 647 mouse IgG1,κ isotype control (BD Pharmingen), and Hoechst 33258 (Thermo Fisher Scientific) and analyzed on a FACSAria instrument (BD).

## Results

### Tofacitinib targets the BM microenvironment and reverses bone marrow stromal cell-mediated growth promotion

To initially validate findings from our drug repurposing screen, we co-cultured the human MM cell line MM.1S, which was included in the screen^8^, with the immortalized bone marrow stromal cell lines HS5 and HS27A. We found that after 24h of co-culture, MM.1S cell numbers approximately doubled compared to monoculture growth after 24h, confirming stromal-induced proliferative signaling in this cell line, as seen in the screen. Tofacitinib treatment for 24 hr reduced MM.1S cell numbers in a dose-dependent manner, such that at >1 μM tofacitinib, MM.1S cell numbers in coculture return to approximately monoculture levels (Fig. 1A). Tofacitinib has no effect on MM.1S cell viability alone nor on HS5 or HS27A alone (Fig. 1B). We further studied the effect of tofacitinib on several other immortalized MM cell lines. In monoculture we found that tofacitinib only demonstrates anti-MM activity in the IL-6 secreting line U266 and in the IL-6 dependent cell line INA-6, with minimal to no effect on the other MM cell lines (Fig. 1C). We further evaluated four myeloma cell lines (U266, MM.1S, RPMI-8226, JJN-3) in which luciferase was stably expressed, allowing for distinction of MM cell viability versus stromal cell viability in co-culture, in the compartment-specific bioluminescence assay^21^. After 24 hr of growth followed by 24 hr of drug exposure, only the stromal-responsive cell lines MM.1S and U266 exhibit any sensitivity to tofacitinib treatment (Fig. 1D). Taken together, these results suggest that tofacitinib selectively targets the growth-promoting interaction between MM cells and the stromal microenvironment. These results further suggest that MM.1S is an appropriate cell line candidate for further study, as it recapitulates the growth-promoting effects of the bone marrow microenvironment known to occur in patients.

**Figure 1.**
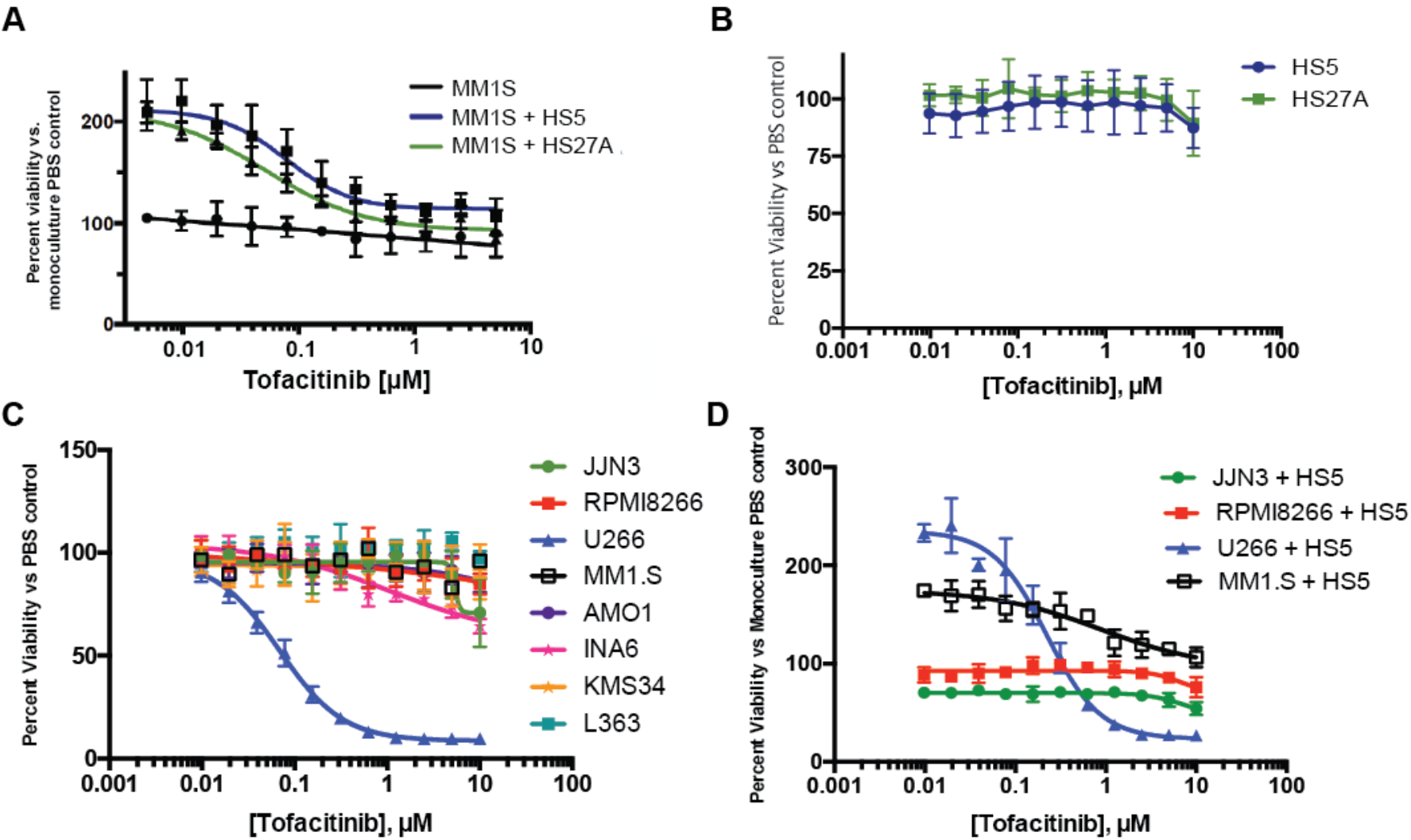
Tofacitinib inhibits stromal-cell proliferation in MM cells mediated by IL-6. A. Tofacitinib has no effect vs. MM1S MM cells in monoculture but instead reverses proliferation induced by bone marrow stromal cell lines HS5 and HS27A. **B.** Tofacitinib has no viability effect vs. bone marrow stromal cells. **C.** Tofacitinib has minimal effects vs. most MM cell lines in mono-culture, except the IL-6 secreting line U266 and IL-6 dependent line INA-6. **D.** In stromal cell co-culture, tofacitinib does not have anti-MM effects vs. JJN-3 and RPMI-8226 cell lines, which do not proliferate in response to stroma. All error bars represent +/-S.D. from assay performed in quadruplicate in 384-well plates.

### Tofacitinib inhibits IL-6 driven growth promotion in coculture with bone marrow stromal cells

To further characterize the nature of pro-growth signaling between the stromal cell microenvironment and MM plasma cells, we performed RNA-seq on MM.1S cells grown alone or grown in co-culture with HS5 stromal cells. We first noticed that the most significantly upregulated transcript in MM.1S in the co-culture setting was SOCS3, part of a well-characterized negative feedback mechanism strongly induced by JAK-STAT activation^10^ (Fig. 2A). We further examined all 67 transcripts significantly upregulated in MM.1S in the co-culture vs. monoculture setting (*p* < 0.005 per DESeq tool^24^; listed in Supplementary Dataset 1). Using the Enrichr tool^25^, ChEA analysis of ChIP-seq datasets^28^ found the most significant enrichment of STAT3-binding sites at the promoter of these upregulated transcripts, among all transcription factors (Fig. 2B). Furthermore, Panther^29^ pathway analysis found the only two significantly enriched pathways to be related to JAK/STAT signaling and interleukin signaling (Fig. 2C). Taken together, these RNA-seq findings are most consistent with a mechanism where IL-6 secreted from stromal cells mediates proliferation by activating JAK/STAT signaling, with STAT3 playing a central role^30^.

**Figure 2.**
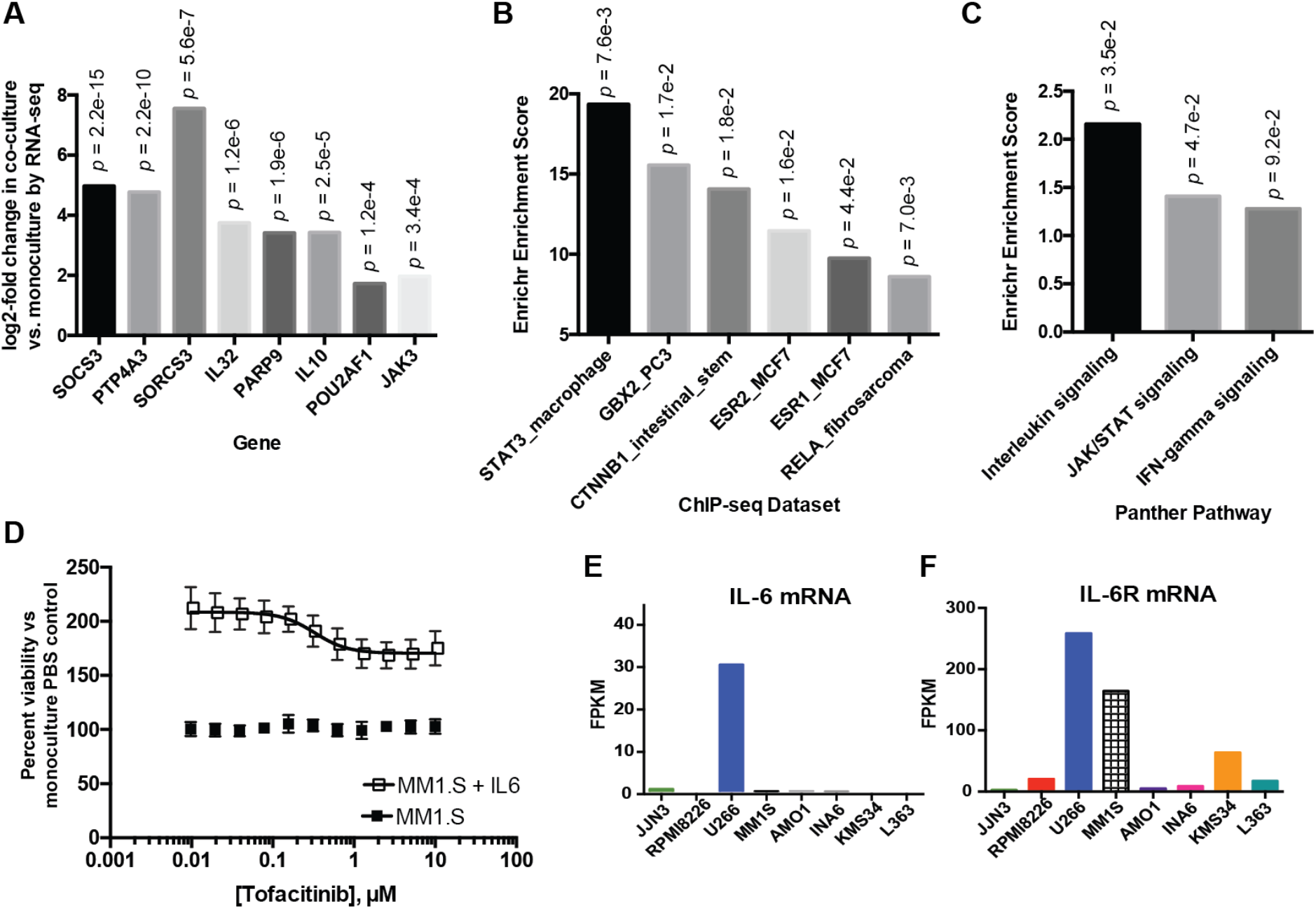
Stromal-induced signatures in MM.1S identified by transcriptome analysis. **A.** Examples of significantly upregulated in genes in MM.1S cells co-cultured with HS5 stromal cells in comparison to MM.1S grown in monoculture. **B.** ChEA analysis of 72 significantly upregulated transcripts from untreated MM.1S in HS5 co-culture vs. monoculture (*p* < 0.05 based on DESeq analysis) demonstrates a significant enrichment of STAT3 transcription-factor binding sites based on ChIP-seq data. **C.** Panther pathway analysis of this gene list demonstrates significant upregulation of interleukin signaling and JAK-STAT signaling among. **D.** Tofacitinib reverses proliferation induced by recombinant IL-6 in MM.1S (50 ng/mL). Error bars represent +/-S.D. from assay performed in quadruplicate in 384-well plates. **E.-F.** Analysis of cross-cell-line RNA-seq data at http://www.keatslab.org/data-repository demonstrates a potential relationship between IL-6 and/or IL-6R gene expression and sensitivity to tofacitinib.

We further investigated this mechanism by confirming that MM.1S cells stimulated with 50 ng/mL recombinant human IL-6 exhibited growth promotion similar to that induced by coculture with stromal cells, suggesting that IL-6 is the primary cytokine mediating stromal-cell induced proliferation (Fig. 2D). Tofacitinib was capable of partially reversing this IL-6 induced growth promotion in MM.1S at drug concentrations similar to those found for phenotypic effects for stromal cell co-culture. We note that we did not find complete reversal of the growth promotion, however, perhaps indicating that the high doses of recombinant IL-6 used here may lead to residual proliferative signaling not fully blocked by tofacitinib.

We further took advantage of publicly available RNA-seq data across a large panel of MM cell lines (http://www.keatslab.org/data-repository) to look for relationships between IL-6 and IL-6R expression and sensitivity to tofacitinib (Fig. 3E-F). Consistent with prior studies^31^, only U266 expresses detectable levels of IL-6, whereas both U266 and MM.1S express relatively high levels of IL-6R compared to other cell lines examined in Fig. 1, suggesting a relationship between IL-6, IL-6R, and sensitivity to tofacitinib.

**Figure 3.**
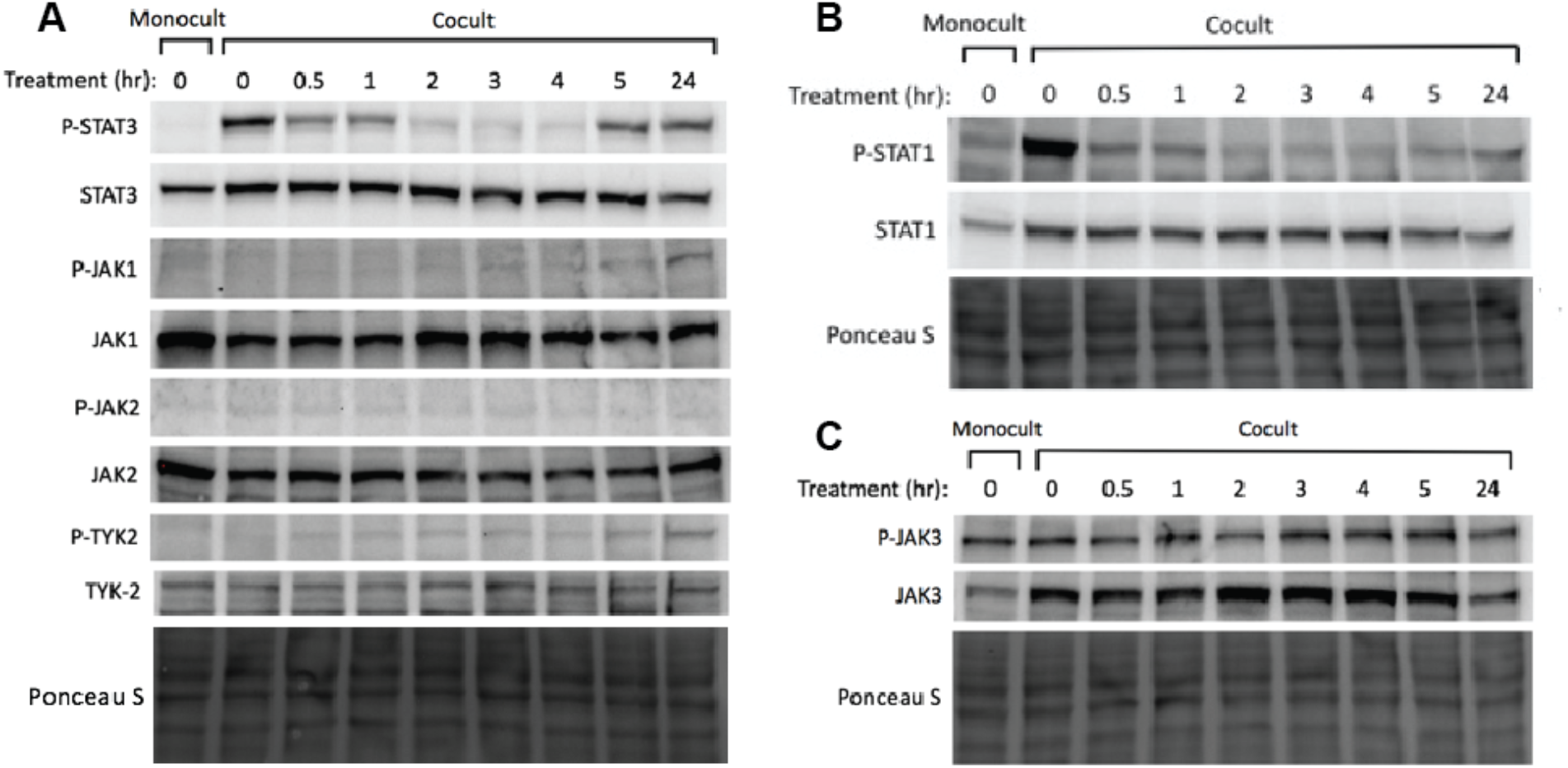
Tofacitinib inhibits JAK/STAT signaling in MM cells in a time-dependent manner. **A.** Western blotting demonstrates a marked increase in STAT3 phosphorylation in untreated co-culture vs. monoculture in MM.1S cells. After treatment with 1 μM tofacitinib there is a rapid decrease in STAT3 phosphorylation with rebound by 24 hr. **B.** STAT1 phosphorylation is also increased in response to co-culture and rapidly reversed by tofacitinib treatment. **C.** While co-culture increases JAK3 protein expression, there does not seem to be any significant change in signaling via JAK3 after tofacitinib based on phosphorylation status. Western blots are representative of assays performed in biological duplicate.

### Tofacitinib inhibits JAK/STAT signaling

Given the proposed mechanism above, we chose to further evaluate downstream effects of IL-6-mediated stimulation and subsequent tofacitinib inhibition of the JAK/STAT pathway. Two of the primary downstream mediators of IL-6R and JAK activation are thought to be pro-proliferation signaling by STAT3 and inhibitory signaling by STAT1^30^. As expected, we found that STAT3 and STAT1 phosphorylation in MM.1S dramatically increases when in coculture with HS5 (Fig. 3A and 3B). 1 μM tofacitinib inhibits STAT3 phosphorylation in MM.1S cells in coculture almost to monoculture level by 2 hrs of treatment, suggestive of on-target effects. Phosphorylation of JAK1, JAK2, and TYK2, which can also activate STAT3, were also studied. We found evidence of a “rebound” effect by 24h of treatment, mediated by well-characterized feedback mechanisms^10^, serving to phosphorylate the JAKs and subsequently re-activate STAT3. Despite this rebound of STAT3 activation, however, MM growth continues to be inhibited based on dose-response results of tofacitinib treatment at 24 hours.

As tofacitinib is known to potently inhibit JAK3, also of interest was an increase in JAK3 expression in MM.1S in coculture versus monoculture, found both by RNA-seq (Fig. 2A) and Western blot (Fig. 3C). However, tofacitinib did not lead to any significant decrease in JAK3 phosphorylation (Fig. 3C). We also evaluated signaling through JAK3’s primary pro-proliferative downstream mediator STAT5 (ref. 32). We found no evidence of STAT5 phosphorylation in either monoculture or co-culture (Supplementary Fig. 2A). These results suggest that JAK3/STAT5 signaling is less central to stroma-supported MM growth. We further confirmed this result using JAK3i, a newly-described, highly-specific, irreversible JAK3 inhibitor^32^. JAK3i had no effect on MM.1S in mono- or coculture (Fig. 4A), nor did it show any synergy with carfilzomib treatment (Supplementary Figure 2B-C).

**Figure 4.**
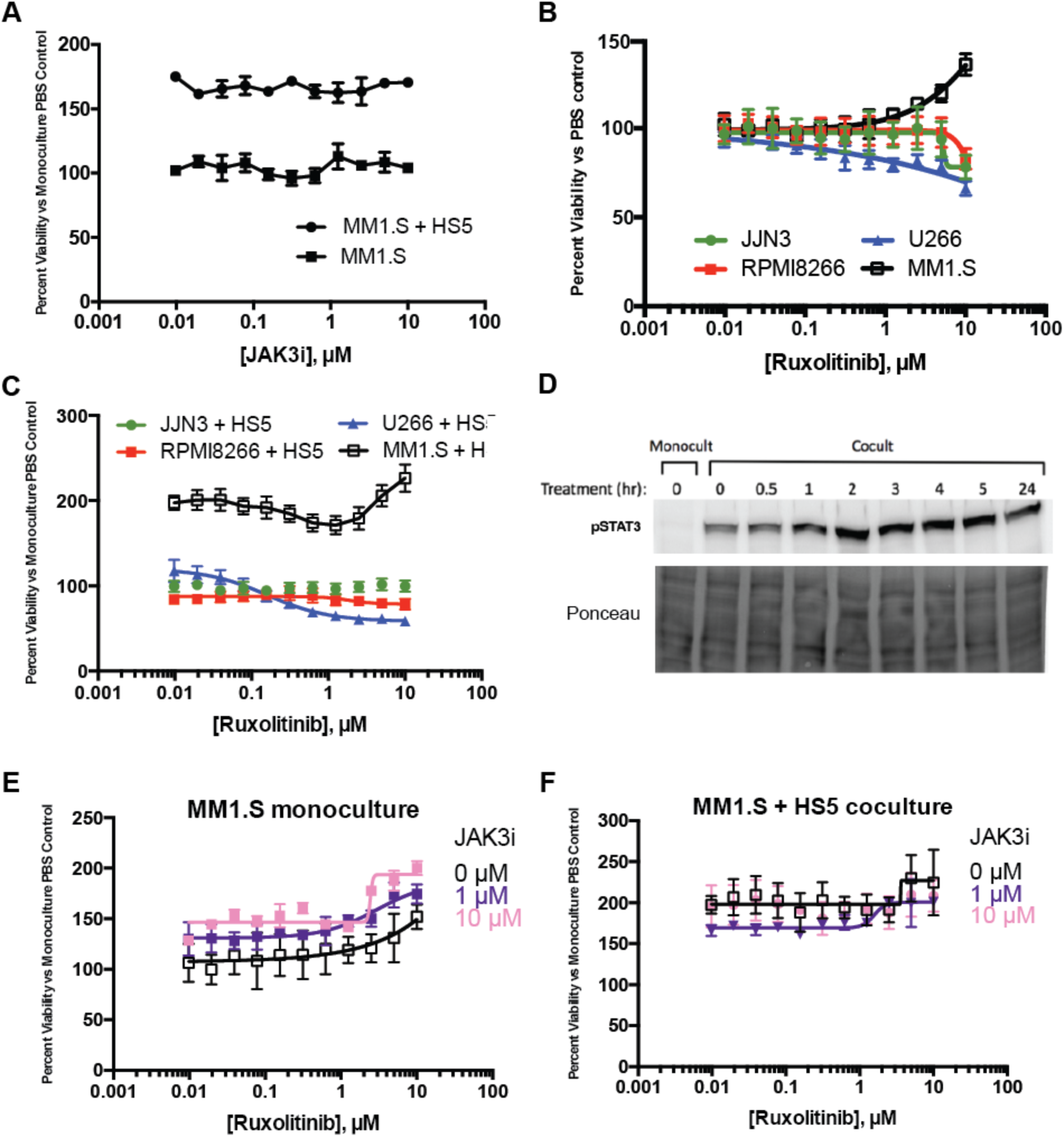
Ruxolitinib demonstrates less anti-MM activity than tofacitinib. **A.** A highly selective, irreversible inhibitor of JAK3, JAK3i, does not have any effects on MM.1S either in mono-culture or in co-culture with HS5. **B.** The JAK1/2 inhibitor ruxolitinib has minimal anti-MM effects in monoculture vs. four MM cell lines, and in fact appears to promote growth of MM.1S at higher concentrations. **C.** A similar phenomenon is noted in HS5 co-culture. **D.** Ruxolitinib does not inhibit and in fact increases signaling via STAT3 in MM.1S grown in HS5 co-culture. **E.-F.** Combination of ruxolitinib with JAK3i, to achieve simultaneous JAK1/2/3 inhibition, does not recapitulate effects of tofacitinib in MM.1S. All error bars represent +/-S.D. from assay performed in quadruplicate in 384-well plates.

Taken together, these results suggest a mechanism whereby stromal cell-induced MM proliferation is mediated through STAT3 transcriptional effects. These signaling pathways are inhibited in an on-target fashion by tofacitinib, ultimately leading to reversal of the proliferation phenotype.

### Ruxolinitib has less anti-MM activity than tofacitinib

Given that our results above suggest a more important role for JAK1 and/or JAK2 in stromal-induced MM proliferation than JAK3, we turned to an alternate candidate for drug repurposing, ruxolinitib. This agent, sold under the trade name Jakofi, is FDA-approved for use in myeloproliferative neoplasms and has much higher affinity for JAK1 and JAK2 over JAK3 or TYK2 (ref. 33). Treatment of U266 cells showed some anti-myeloma effect (Fig. 4B-C) but this was significantly reduced compared to tofacitinib (Fig. 1). Surprisingly, treatment of MM.1S actually showed a promotion of growth at higher concentrations, both in the monoculture and co-culture settings (Fig. 4B-C). Western blotting demonstrated that 1 μM ruxolitinib was unable to inhibit STAT3 activation in MM.1S in co-culture (Fig. 4D). In fact, STAT3 phosphorylation increased after 2 hr of treatment, consistent with the pro-proliferative effect seen in Fig. 4B-C. To confirm that these effects could not be rescued with simultaneous inhibition of JAK1/2/3, as accomplished by tofacitinib, we combined ruxolitinib with JAK3i. We found this combination was also insufficient to recapitulate tofacitinib’s effects in mono- or co-culture (Fig. 4E-F). Taken together, these findings suggest that ruxolinitib is unable to inhibit pro-proliferative STAT3 signaling in an in MM.1S cells, thereby supporting tofacitinib as having greater potential as a repurposed anti-myeloma therapy.

### Unbiased phosphoproteomics demonstrates that tofacitinib broadly reverses pro-growth signaling induced by bone marrow stroma

Our targeted investigations above specifically focused on the JAK/STAT pathway. To further elucidate the mechanism of tofacitinib in this system, as well as gain a broader view of stromal-induced proliferation and signaling in MM cells, we pursued unbiased mass spectrometry (MS)-based phosphoproteomics. We studied four samples, performed in biological replicate: 1) MM.1S cells in monoculture; MM.1S cells in co-culture, either 2) untreated (DMSO control); 3) treated with 1 μM tofacitinib for 1.5 hr; or 4) treated with 1 μM tofacinitib for 24 hr. We harvested MM.1S cells in suspension, enriched for phosphorylated peptides using immobilized metal chromatography, and analyzed by LC/MS-MS with peptide quantification performed using MaxQuant^27^.

In total 4862 phosphopeptides had intensity data in all four samples and were used for further analysis (listed in Supplementary Dataset 2). >99% of these sites are serine and threonine phosphorylation events, consistent with other phosphoproteomic studies using this enrichment method^34^. We first evaluated for phosphosites with >4-fold intensity increases in untreated co-culture vs. monoculture. Using our RNA-seq data as a proxy for protein-level changes, we verified these phospho-site changes were largely driven by changes in signaling and not protein abundance (Supplementary Fig. 3). Panther pathway analysis revealed the only significantly enriched pathway among the 544 upregulated phosphosites to be JAK/STAT signaling (Fig. 5A). However, based on Kinase Enrichment Analysis^35^, we found enriched signatures not only of JAK1 substrates, but also other kinases driving proliferation and the cell cycle such as mTOR, CDK1, and CDK2 (Fig. 5B). These findings demonstrate that unbiased phosphoproteomics can uncover broad signaling effects of the bone marrow microenvironment even downstream of JAK/STAT.

**Figure 5.**
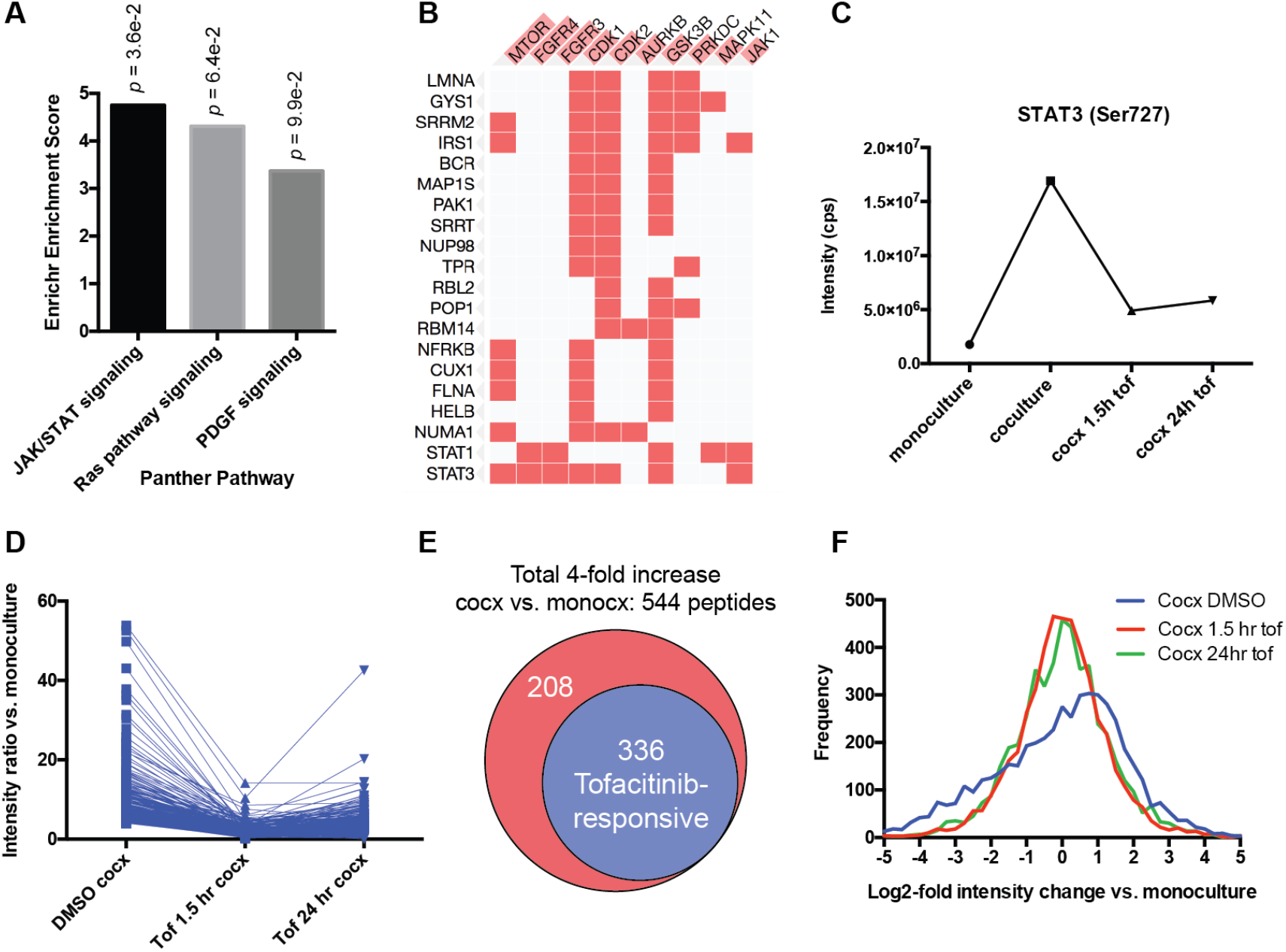
Unbiased phosphoproteomics reveals that tofacitinib broadly reverses pro-growth signaling from stroma to MM cells. **A.** Analysis of 4895 phosphorylation sites quantified by LC-MS/MS on biological replicate samples revealed 544 phosphosites to be upregulated 4-fold in untreated MM.1S in coculture with HS5 vs. monoculture. Of these upregulated phosphopeptides, Panther pathway analysis showed JAK/STAT signaling to be the only significantly enriched pathway (*p* = 0.036). **B.** Kinase Enrichment Analysis demonstrated that many of the upregulated phosphopeptides derive from known substrates of other kinases related to proliferation such as mTOR, CDK1, and CDK2, as well as JAK1. Phosphorylated protein substrates are on the left and enriched kinases across the top. Length of red bar in kinase name is indicative of strength of enrichment. **C.** Quantitative phosphoproteomic intensity of the JAK-responsive phosphosite Ser727 on STAT3, both in untreated co-culture vs. monoculture, and after tofacitinib treatment, is very similar to the pattern found by Western blotting for the other known-JAK responsive STAT3 phosphosite Tyr707 (Fig. 3A), serving to validate this proteomics approach. **D.** Dynamics of all phosophosites found to be responsive to tofacitinib based on the criteria: 1) increased 4-fold in co-culture vs. monoculture 2) decreased at least-2 fold from untreated co-culture after 1.5 hr of 1 μM tofacitinib treatment and 3) phosphosite intensity remains below the untreated co-culture level after 24h of tofacitinib treatment. **E.** Of 544 upregulated phosphopeptides in co-culture, 336 (62%) were defined as being tofacitinib-responsive using the criteria in D. **F.** Examining the global phosphosite intensity across all 4895 quantified phosphopeptides demonstrates a general increase in phosphorylation of MM.1S proteins in untreated co-culture with HS5 (“Cocx DMSO”; note shift of distribution maximum to log2-fold change vs. monoculture of ~1) which is then broadly reversed by tofacitinib treatment. Tof = tofacitinib; monocx = monoculture; cocx = co-culture.

Next, for validation of effects of tofacitinib treatment, we first examined Ser727 on STAT3 (Fig. 5C), a known JAK-responsive phosphosite^30^. Our quantitative MS results were remarkably in line with Western blotting for another JAK-responsive phosphosite on STAT3, Tyr707 (Fig. 3A), with a very large increase in both phosphosites in untreated co-culture compared to baseline, a >2-fold decrease in phosphorylation after short-term tofacitinib treatment, and a rebound in phosphorylation at 24 hr.

This finding both serves to validate our phosphoproteomic data as well as help us define a signature of tofacitinib-responsive phosphosites. Remarkably, 336 of the 544 up-regulated phosphosites in untreated co-culture vs. monoculture (62%) met the same criteria of being tofacitinib-responsive (Fig. 5D-E). Furthermore, examination across all measured phosphosites demonstrates that while phosphorylation is broadly increased in the co-culture setting, noted as a general shift toward positive phosphosite intensities in the untreated sample, treatment with tofacitinib largely reverses this finding, re-creating a normal distribution around the intensity values found in monoculture (Fig. 5F). Taken together, these findings demonstrate that tofacitinib broadly reverses the signaling pathways driving stromal-incuded proliferation in MM cells, both at the level of direct JAK targets as well downstream proliferative signals, informing our mechanistic understanding of this treatment beyond targeted Western blots alone.

### Tofacitinib does not appear to have significant off-target activity

Another advantage of unbiased phosphoproteomics is potentially detecting additional off-target effects mediating tofacitinib response. We filtered for peptides that appeared unaffected by stromal-induced signaling (less than +/-50% intensity change in untreated co-culture vs. monoculture) that decreased in intensity >4-fold after 1.5 hr tofacitinib treatment (Fig. 6A). We identified only 54 peptides that fit this filter, and neither Panther nor Kinase Enrichment Analysis identified any enriched signatures (not shown). Given the small number of peptides and lack of any biological signatures, it appears most likely these 54 peptides are background noise in the data and do not suggest any significant off-target effects of tofacitinib.

**Figure 6.**
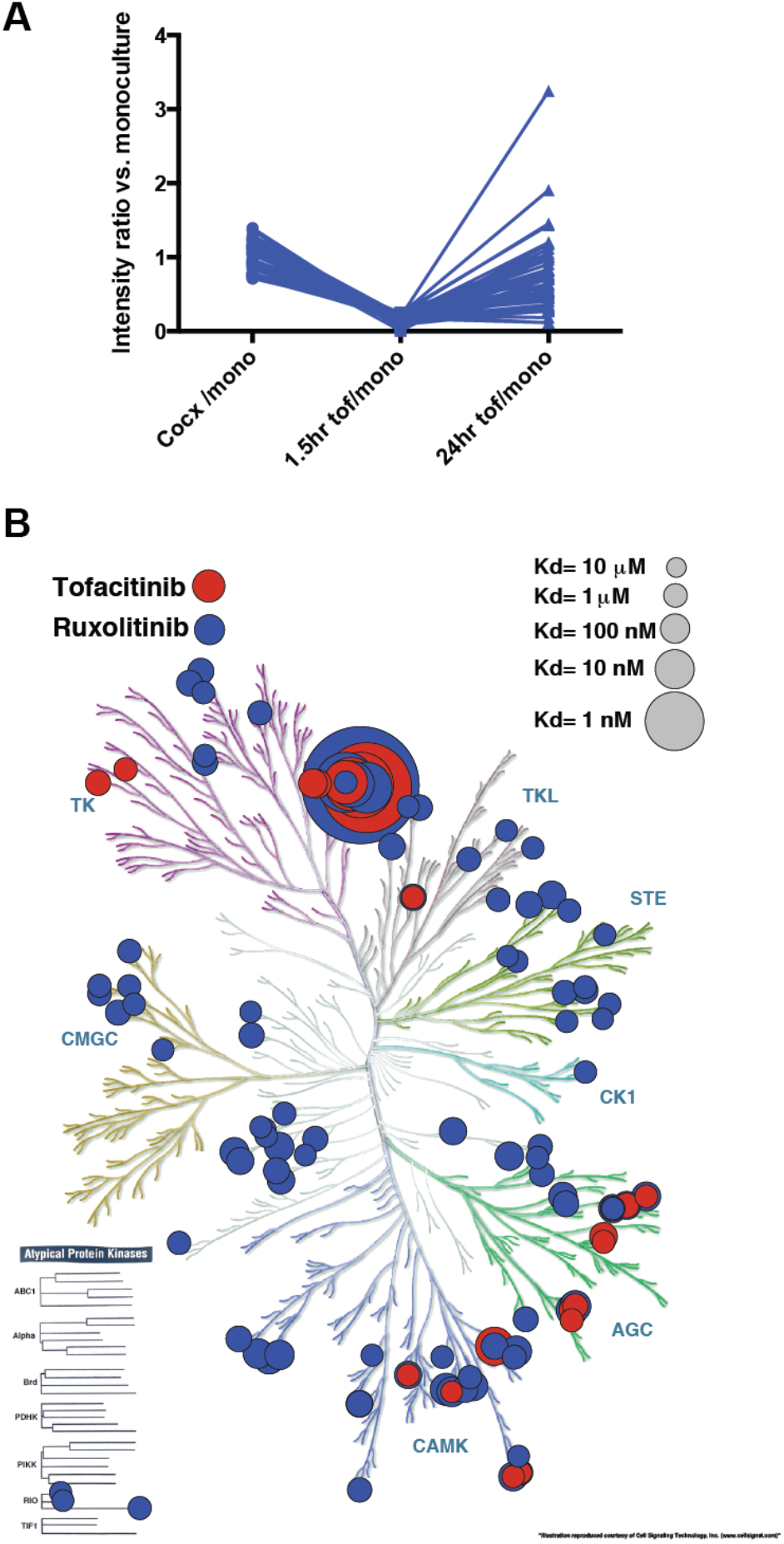
Phosphoproteomics reveals minimal evidence of tofacitinib off-target effects in MM. **A.** Only 54 phosphopeptides were identified that met the criteria: 1) <50% intensity change in untreated co-culture vs. monoculture and 2) decreased >4-fold after 1.5 hr tofacitinib treatment. These phosphopeptides showed no significant biological pathway enrichment upon Panther or other bioinformatic analysis by Enrichr, suggesting no prominent off-target effects of tofacitinib in this system. **B.** Cell-free kinase inhibitor activity data from LINCS KINOMEscan database demonstrates that tofacitinib is much more specific for JAK-family kinases (large circles, denoting strong binding, top center) than ruxolitinib, which has off-target activity against many more kinases. Figure rendered using KinomeRender^39^.

However, to further investigate the possibility of any off-target effects of tofacitinib, we downloaded available data from the LINCS KINOMEscan database (http://lincs.hms.harvard.edu/kinomescan/) and plotted versus the human kinase phylogenetic tree (Fig. 6B). These results demonstrate that, at least in cell-free assays, tofacinitib shows much greater specificity for JAK-family kinases compared to ruxolitinib, which has numerous off-target activities. These findings underscore that tofacinitib’s effects in this co-culture model appear to be due to on-target activity. Furthermore, we speculate that through an as-yet-undefined mechanism, off-target effects of ruxolitinib may antagonize on-target JAK/STAT inhibition in these MM models, leading to decreased efficacy of ruxolitinib compared to tofacitinib.

### Tofacitinib has anti-MM activity in the bone marrow microenvironment *in vivo*

Toward the goal of repurposing tofacitinib as an anti-MM therapy in patients, we next examined the efficacy of tofacitinib *in vivo*. For this orthotopic disseminated xenograft model we used luciferase-labeled MM.1S and U266 cell lines, which specifically home to the murine bone marrow after intravenous implantation in NOD *scid* gamma (NSG) mice. Treatment was initiated after two weeks of tumor growth and continued for four weeks at ~2/3 of the maximal tolerated dose of tofacitinib (21.5 mg/kg/day by subcutaneous infusion)^36^. Encouragingly, we found significantly increased murine survival in both of these cell line models (Fig. 7A-B) as well as decreased tumor burden based on bioluminescent imaging quantification (Fig. 7C-D). These findings indicate that despite the *in vitro* rebound we observed in STAT3 activation (Fig. 3A), continuous treatment with tofacitinib still leads to anti-myeloma effects *in vivo*.

**Figure 7.**
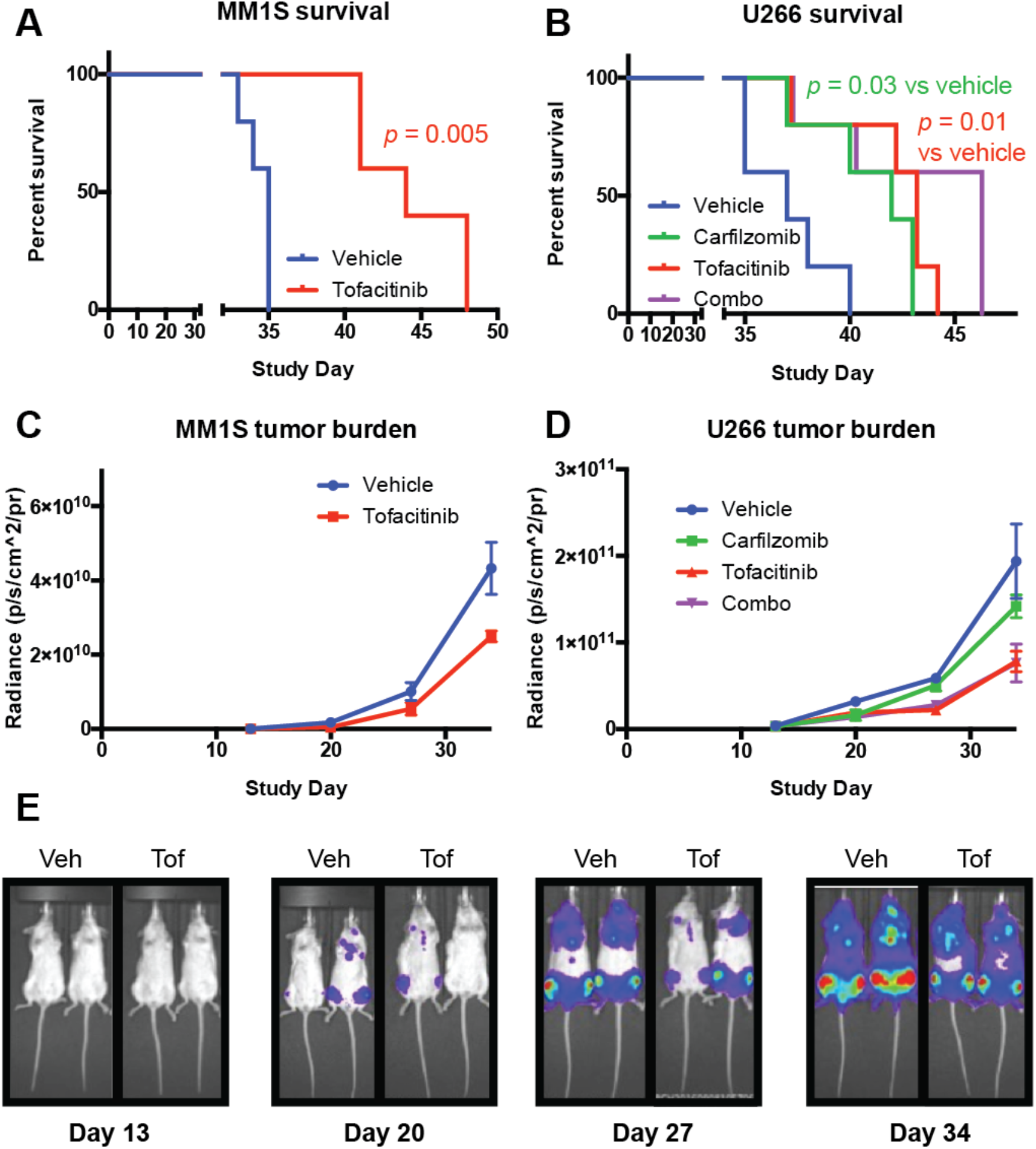
Tofacitinib has anti-MM activity *in vivo*. Luciferase labeled-MM.1S and U266 MM cell lines were implanted intravenously into NSG mice and tumors allowed to grow for 13 days. Mice were randomized (*n* = 5 mice per arm) and drug dosing was then begun for four weeks. Tofacitinib dose = 21.5 mg/kg/day by subcutaneous pump. Carfilzomib dose = 2 mg/kg IV 2x/week for four weeks. **A-B.** Tofacitinib significantly increased survival of NSG mice in these aggressive mouse models of MM by Log-rank test. **C-D.** Bioluminescence imaging for tumor burden was performed at Day 13, Day 20, Day 27, Day 34. Error bars represent +/-S.D. **E.** Example bioluminescent images from MM.1S study showing prominent localization of tumor cells to hindlimb bone marrow.

We also performed *in vitro* studies of tofacitinib in combination with first-line myeloma therapies carfilzomib and melphalan (Supplementary Fig. 4). We found combination effects with both of these agents in the co-culture model. Based on these results, in our U266 murine study we also included study arms with carfilzomib alone at 2/3 of the maximal tolerated dose (2 mg/kg IV, 2x/week) and a combination of carfilzomib and tofacitinib at the above doses. We found that tofacitinib led to similar lifespan extension as carfilzomib monotherapy, and also found evidence of additive effects of the two agents, though the difference between the combination and either monotherapy was not statistically significant.

We also tested tofacitinib versus two primary MM bone marrow samples treated *ex vivo*. We did not see any significant viability effects of tofacitinib in either of these samples, against either malignant plasma cells or other normal bone marrow mononuclear cells (Supplementary Fig. 5). However, this result appears consistent with our *in vitro* and phosphoproteomic results, which suggest that IL-6 dependent plasma cell proliferation is necessary for tofacitinib to have any effect. Primary MM plasma cells isolated *ex vivo* in 2D culture are known to have minimal ability to actively proliferate even in the presence of cytokines or stromal stimulation^37^. Therefore, these results may more reveal the limitations of *ex vivo* patient assays in MM rather than preclude the therapeutic efficacy of tofacitinib in MM patients, where plasma cells are constantly proliferating in an IL-6-dependent manner within the bone marrow.

## Discussion

Given the importance of the bone marrow microenvironment for MM pathogenesis, the IL-6/JAK/STAT signaling axis has generated significant interest as a therapeutic target in MM. JAK inhibition has already been validated in a number of preclinical studies as a way to target this pathway^15–20^. However, the studied compounds are not yet available clinically and may never be. Here, following the results of a large scale repurposing screen, we validated tofacitinib as a potential therapy that can be rapidly translated into MM patients.

Using a combination of mechanistic pharmacology and unbiased mass spectrometry-based phosphoproteomics, we found that tofacitinib appears to inhibit stromal-induced proliferation of MM plasma cells by inhibiting the IL-6/JAK/STAT signaling axis. Our results support the use of unbiased phosphoproteomics both in kinase inhibitor evaluation and more broadly in MM biology, where this method has been only applied in a very limited fashion.

Surprisingly, in our studies we found that the FDA-approved JAK1/2 inhibitor ruxolitinib did not lead to the same anti-myeloma effects as tofacitinib, and in fact may paradoxically promote plasma cell proliferation at high concentrations. We do note that ruxolitinib was previously evaluated in a small trial of 13 MM patients in combination with dexamethasone (NCT00639002) and no significant anti-MM effects were noted in this small study. Our *in vitro* results here may provide a partial explanation for the lack of ruxolitinib activity in that trial.

We note that these studies are of course limited in that they are performed in MM cell lines. While we primarily focused our analysis on the stromal-responsive cell lines MM.1S and U266, which appear to better mimic the malignant plasma cell phenotype found in MM patients, at this point focused clinical trials in MM will be necessary to truly evaluate whether tofacitinib has anti-MM effects.

Toward this goal, the value of drug repurposing becomes readily apparent. Tofacitinib can be quickly moved into Phase I/II studies in MM as the tolerated doses and adverse event profiles of this drug are well-characterized in humans^11^. Tofacitinib is generally well-tolerated at therapeutic doses in RA^11^ and, despite early concerns, post-market surveillance studies have not revealed any increased risk of malignancies compared to other RA therapies^38^. Intriguingly, a patient population with both MM and RA could readily serve as the basis of a multi-center trial in combination with standard of care MM therapies. Alternatively, a patient population with early-stage disease, perhaps smoldering myeloma, may be the optimal setting for clinical use, when plasma cells may be most dependent on IL-6 and other microenvironment cues. In conclusion, tofacitinib is a promising agent to reverse the tumor-proliferative effects of the bone marrow microenvironment that can be rapidly repurposed to benefit MM patients.

## Acknowledgements

This work was supported by the UCSF Stephen and Nancy Grand Multiple Myeloma Translational Initiative and The Myeloma Research Fund of the Silicon Valley Community Foundation (to B.T.A. and A.P.W.), and an NCI Clinical Scientist Development Award (K08 CA184116), a Dale F. Frey Breakthrough Award from the Damon Runyon Cancer Research Foundation (DFS 14-15), and an American Cancer Society Individual Research Award (IRG-97-150-13) (to A.P.W.). We thank Drs. Jeffrey Wolf, Tom Martin, Nina Shah, and Cammie Edwards for discussions, advice, and insight. We thank the staff of the UCSF Helen Diller Family Cancer Center Preclinical Therapeutic Core facility, supported by NCI Cancer Center Support Grant P30 CA082103, for completion of murine studies. We thank Dr. Diego Acosta-Alvear for providing luciferase-labeled MM cell lines.

## REFERENCES

1 Manier, S., Kawano, Y., Bianchi, G., Roccaro, A. M. & Ghobrial, I. M. Cell autonomous and microenvironmental regulation of tumor progression in precursor states of multiple myeloma. Curr Opin Hematol 23, 426–433 (2016).

2 Kuehl, W. M. & Bergsagel, P. L. Molecular pathogenesis of multiple myeloma and its premalignant precursor. J Clin Invest 122, 3456–3463 (2012).

3 Bianchi, G. & Munshi, N. C. Pathogenesis beyond the cancer clone(s) in multiple myeloma. Blood 125, 3049–3058 (2015).

4 Gupta, S. C., Sung, B., Prasad, S., Webb, L. J. & Aggarwal, B. B. Cancer drug discovery by repurposing: teaching new tricks to old dogs. Trends Pharmacol Sci 34, 508–517 (2013).

5 Corsello, S. M. et al. The Drug Repurposing Hub: a next-generation drug library and information resource. Nat Med 23, 405–408 (2017).

6 Shim, J. S. & Liu, J. O. Recent advances in drug repositioning for the discovery of new anticancer drugs. Int J Biol Sci 10, 654–663 (2014).

7 Nosengo, N. Can you teach old drugs new tricks? Nature 534, 314–316 (2016).

8 Murnane, M. et al. Defining Primary Marrow Microenvironment-Induced Synthetic Lethality and Resistance for 2,684 Approved Drugs Across Molecularly Distinct Forms of Multiple Myeloma. Blood (ASH Abstract) 126 (2015).

9 Meyer, D. M. et al. Anti-inflammatory activity and neutrophil reductions mediated by the JAK1/JAK3 inhibitor, CP-690,550, in rat adjuvant-induced arthritis. J Inflamm 7, 41 (2010).

10 Shuai, K. & Liu, B. Regulation of JAK-STAT signalling in the immune system. Nature reviews. Immunology 3, 900–911 (2003).

11 Fleischmann, R. et al. Placebo-controlled trial of tofacitinib monotherapy in rheumatoid arthritis. N Engl J Med 367, 495–507 (2012).

12 Klein, B., Zhang, X. G., Lu, Z. Y. & Bataille, R. Interleukin-6 in human multiple myeloma. Blood 85, 863–872 (1995).

13 Das, R. et al. Microenvironment-dependent growth of preneoplastic and malignant plasma cells in humanized mice. Nat Med 22, 1351–1357 (2016).

14 Rosean, T. R. et al. Preclinical validation of interleukin 6 as a therapeutic target in multiple myeloma. Immunol Res 59, 188–202 (2014).

15 Monaghan, K. A., Khong, T., Burns, C. J. & Spencer, A. The novel JAK inhibitor CYT387 suppresses multiple signalling pathways, prevents proliferation and induces apoptosis in phenotypically diverse myeloma cells. Leukemia 25, 1891–1899 (2011).

16 Li, J. et al. INCB16562, a JAK1/2 selective inhibitor, is efficacious against multiple myeloma cells and reverses the protective effects of cytokine and stromal cell support. Neoplasia 12, 28–38 (2010).

17 De Vos, J., Jourdan, M., Tarte, K., Jasmin, C. & Klein, B. JAK2 tyrosine kinase inhibitor tyrphostin AG490 downregulates the mitogen-activated protein kinase (MAPK) and signal transducer and activator of transcription (STAT) pathways and induces apoptosis in myeloma cells. Brit J Hematol 109, 823–828 (2000).

18 Ramakrishnan, V. et al. TG101209, a novel JAK2 inhibitor, has significant in vitro activity in multiple myeloma and displays preferential cytotoxicity for CD45+ myeloma cells. Am J Hematol 85, 675–686 (2010).

19 Scuto, A. et al. The novel JAK inhibitor AZD1480 blocks STAT3 and FGFR3 signaling, resulting in suppression of human myeloma cell growth and survival. Leukemia 25, 538–550 (2011).

20 Burger, R. et al. Janus kinase inhibitor INCB20 has antiproliferative and apoptotic effects on human myeloma cells in vitro and in vivo. Mol Cancer Therap 8, 26–35 (2009).

21 McMillin, D. W. et al. Tumor cell-specific bioluminescence platform to identify stroma-induced changes to anticancer drug activity. Nat Med 16, 483–489 (2010).

22 Smith, E. J. et al. A novel, native-format bispecific antibody triggering T-cell killing of B-cells is robustly active in mouse tumor models and cynomolgus monkeys. Sci Rep 5, 17943 (2015).

23 Wiita, A. P. et al. Global cellular response to chemotherapy-induced apoptosis. eLife 2, e01236 (2013).

24 Anders, S. & Huber, W. Differential expression analysis for sequence count data. Genome Biol 11, R106 (2010).

25 Kuleshov, M. V. et al. Enrichr: a comprehensive gene set enrichment analysis web server 2016 update. Nucleic Acids Res 44, W90–97 (2016).

26 Fila, J. & Honys, D. Enrichment techniques employed in phosphoproteomics. Amino Acids 43, 1025–1047 (2012).

27 Cox, J. & Mann, M. MaxQuant enables high peptide identification rates, individualized p.p.b.-range mass accuracies and proteome-wide protein quantification. Nat Biotechnol 26 (2008).

28 Lachmann, A. et al. ChEA: transcription factor regulation inferred from integrating genome-wide ChIP-X experiments. Bioinformatics 26, 2438–2444 (2010).

29 Mi, H. et al. PANTHER version 11: expanded annotation data from Gene Ontology and Reactome pathways, and data analysis tool enhancements. Nucleic Acids Res 45, D183-D189 (2017).

30 Nelson, E. A., Walker, S. R. & Frank, D. A. in Advances in Biology and Therapy of Multiple Myeloma: Volume 1: Basic Science (ed N.C. Munshi and K.C. Anderson) Ch. 7, 117–138 (Springer, 2013).

31 Jernberg-Wiklund, H., Pettersson, M., Carlsson, M. & Nilsson, K. Increase in interleukin 6 (IL-6) and IL-6 receptor expression in a human multiple myeloma cell line, U-266, during long-term in vitro culture and the development of a possible autocrine IL-6 loop. Leukemia 6, 310–318 (1992).

32 Smith, G. A., Uchida, K., Weiss, A. & Taunton, J. Essential biphasic role for JAK3 catalytic activity in IL-2 receptor signaling. Nat Chem Biol 12, 373–379 (2016).

33 Quintas-Cardama, A. et al. Preclinical characterization of the selective JAK1/2 inhibitor INCB018424: therapeutic implications for the treatment of myeloproliferative neoplasms. Blood 115, 3109–3117 (2010).

34 Leitner, A. Enrichment Strategies in Phosphoproteomics. Methods Mol Biol 1355, 105–121 (2016).

35 Lachmann, A. & Ma’ayan, A. KEA: kinase enrichment analysis. Bioinformatics 25, 684–686 (2009).

36 Yokoyama, S. et al. Janus Kinase Inhibitor Tofacitinib Shows Potent Efficacy in a Mouse Model of Autoimmune Lymphoproliferative Syndrome (ALPS). J Clin Immunol 35, 661–667 (2015).

37 Zlei, M. et al. Characterization of in vitro growth of multiple myeloma cells. Exp Hematol 35, 1550–1561 (2007).

38 Curtis, J. R. et al. Tofacitinib, an oral Janus kinase inhibitor: analysis of malignancies across the rheumatoid arthritis clinical development programme. Ann Rheum Dis 75, 831–841 (2016).

39 Chartier, M., Chenard, T., Barker, J. & Najmanovich, R. Kinome Render: a stand-alone and web-accessible tool to annotate the human protein kinome tree. PeerJ 1, e126 (2013).

